# Wnt signaling modulates tissue mechanics, actin order, and regeneration in *Hydra vulgaris*

**DOI:** 10.1101/2025.09.02.673719

**Authors:** Thomas Perros, Stéphane Joly, Adama Mbaye, Philippe Marcq, Olivier Cochet-Escartin

## Abstract

*Hydra vulgaris* is a powerful model organism in the study of axial patterning and regeneration. While recent models emphasize the importance of mechanical cues in establishing the body axis of *Hydra* tissue spheres, the precise role of the Wnt pathway, known to be central to axial patterning, in regulating tissue mechanics and cytoskeletal organization remains unclear. In this paper, we pharmacologically modulated the canonical Wnt pathway using Alsterpaullone and iCRT14 and assessed their effects on regeneration, tissue rheology, osmotic oscillations and actin organization in *Hydra* tissue spheres. We found that Wnt activation prevents full regeneration, softens the constitutive tissues, disrupts osmotic oscillations, leads to an accumulation of dense actin filaments but impairs their orientational order. Conversely, we found that Wnt inhibition partially impairs regeneration and reduces actin filament presence while preserving alignment. These results support the underlying hypotheses of two existing mechano-chemical models of *Hydra* patterning, suggesting that Wnt-driven feedbacks on both tissue stiffness and actin order could contribute to the robust emergence of body axes. Our findings highlight the possibility of redundant and coupled mechanisms in the *Hydra* patterning system, potentially explaining its robustness.

## Introduction

*Hydra vulgaris* is a freshwater cnidarian well-known for its ability for whole body regeneration. Almost any piece of excised tissue, as well as cellular reaggregates can spontaneously reform a full, viable organism in just a few days (1). As such, it has long served as a model organism to study biological patterning and morphogenesis from theoretical and experimental perspectives. With its simple body plan, *Hydra* offers a unique system to study how an organizing axis can arise spontaneously in the absence of external cues. Following the seminal works of Alan Turing on reaction-diffusion systems (2) or of Lewis Wolpert on the French flag model (3), this question was first tackled from a purely biochemical point of view. Gierer and Meinhardt proposed in the 1970s a reaction-diffusion model inspired by Turing’s work and adapted to the specific case of *Hydra* (4). This model reproduced several qualitative features of *Hydra* regeneration and grafting experiments, although successful simulations depended on specific assumptions about tissue size, boundary conditions, and initial activator distributions. It did so through the interplay of two morphogens: a head activator and a head inhibitor. Further, the canonical Wnt pathway in general, and the ligand HyWnt3 in particular were shown to have all the expected properties of the head activator in this model. Not only is this pathway self-activating (5), it was also shown that the Wnt/β-catenin pathway controls the formation of a head organizer (6, 7). Decades of research, however, have failed to uncover an equivalent of the head inhibitor in the Gierer Meinhardt model. Some candidates were identified such as Sp5 (8), Dkk1/2/4 (9), or *Hydra* astacin-7 (10) but none of them have proven to be fully satisfactory for this role, as each showed characteristics that were inconsistent with the predictions of the model. In particular, they did not reproduce the spatial distribution of the inhibitor predicted by the Gierer Meinhardt model. While this did not constitute definitive proof, it suggested that the model remained incomplete.

In recent years, the focus in the field has switched to a mechano-chemical view of *Hydra* regeneration. Key to this change was the observation that osmotically driven mechanical oscillations seemed required for axis definition (11). During their regeneration, excised tissue pieces and cellular re-aggregates start by forming a so-called tissue sphere in which the two main epithelial monolayers, the endoderm and ectoderm, form a spherical lumen. The osmolarity of the inner medium being higher than that of the outside medium, water is actively pumped to the inside of the lumen which starts swelling. It does so to the point where mechanical tension in the tissue becomes too large and the tissues rupture and the lumen quickly deflates. The resulting wound is then healed, and a new cycle begins, leading to a succession of saw-tooth like oscillations. At some point these oscillations show a reduction in amplitude and period, indicative of the formation of a new functional mouth (12) and this has been referred to as a switch from Phase I to Phase II oscillations (11). This showed that the head organizer must be defined prior to this shift in osmotic oscillations. More intriguing was the fact that when these oscillations were blocked by increasing the osmolarity of the outside medium, regeneration failed. Tissue spheres would remain alive but would never show signs of elongation in one direction and no new tentacles (11, 13). This was a clear indication that the formation of the head organizer is, at least partly, mechanically regulated and linked to tissue stretch, tension or rupture. This was later confirmed by molecular analysis which showed a direct feedback loop through which tissue stretch induces the expression of HyWnt3 and other Wnt ligands (13).

In parallel, other researchers have put forward the idea that the organization of supracellular actin filaments, called myonemes, in the ectoderm could be linked to axis definition (14). In the adult *Hydra*, these myonemes align with the organizing axis all over the body column. In an excised tissue piece, this organization can be partially inherited despite the folding back into a spherical shape. This organization seemed to correlate strongly with the axis of the regenerated animal with nematic defects in the organization co-localizing with the future mouth and foot (15). This suggested that spontaneous pattern formation might not be necessary in *Hydra* regeneration from excised tissue pieces. It didn’t explain, however, how patterning is achieved in cellular aggregates which lack any inherited myonemes or why osmotic oscillations would be necessary in the regenerative process.

Nevertheless, both approaches shared an emphasis on mechanical cues in *Hydra* patterning and, to help resolve this question, we recently proposed a mechanical characterization of regenerating *Hydra* tissue spheres using parallelized micro-aspiration experiments (16). We showed that, during osmotic oscillations, they behave as non-linear elastic spherical shells whose mechanical properties were quantified. Based on these different results, two new mechano-chemical models of *Hydra* patterning have emerged in the past year, focusing on different ingredients.

The first model builds upon the results of (14, 15) and highlights the role of actin nematic order in the ectoderm (17). The topology of tissue spheres requires that nematic defects with a total charge of +2 are formed. At the location of a +1 defect, the myonemes were observed to meet in an aster-like configuration. Because of acto-myosin contractility, these defects turn into regions of high tensile stresses and thus high cellular deformations. The model supposes that this local strain triggers a mechano-genetic feedback loop leading to a local activation of the Wnt signaling pathway. To ensure the stabilization of this proto-organizer, the model also hypothesizes that the actin organization responds to a putative Wnt gradient by aligning along its direction, finishing to establish a full nematic order. The fact that an interplay between a morphogen gradient and a nematic field can stabilize nematic defects and drive patterning has been demonstrated theoretically (18), further supporting the rationale of this model. However, this model requires the pre-existence of a partial actin nematic order and thus is not sufficient to explain patterning in regeneration from cellular aggregates where this order is completely lost. It also doesn’t require the existence of osmotic oscillations since cellular deformations result from the contractility of the actin network and not the swelling of the tissue spheres.

The second model focuses on osmotic oscillations and the same hypothesis of a strain-dependent morphogen source (19). By random fluctuations of either deformations or mechanical properties, swelling of the tissue spheres creates a region with slightly larger deformations. In turn, this region starts expressing more of the morphogen. The authors showed that Wnt activation could make the tissue more deformable. Mechanical deformations could thus be locally amplified by a positive feedback loop. In addition, as one region becomes softer and highly deformed, the rest of the tissue sphere is less deformed, limiting the expression of the morphogen. This model thus resembles a classic reaction-diffusion process with a major difference: the long-range inhibitor is not of a chemical nature but of a mechanical one. This exciting idea of a mechano-chemical version of a reaction-diffusion process as a robust patterning system was already studied theoretically in the context of *Hydra* regeneration (20) and embryo development (21). One notable aspect of this second model is that as it does not require any pre-existing actin organization, it could apply for both excised tissue pieces and cellular aggregates. However, it relies on hypotheses and does not require any specific actin nematic order.

To shed light on these apparent contradictions, one needs to test the underlying hypotheses of both models. On the one hand, the effect of tissue deformations on Wnt activation are well established.

On the other, the effects of Wnt activation on rheology and actin organization remain poorly understood. A difficulty in achieving this goal is that, under normal conditions, Wnt activation on *Hydra* tissue spheres is highly localized and difficult to observe *in vivo*. To overcome this limitation, we decided to work on tissue spheres on which the Wnt pathway was either activated or inhibited everywhere.

The aim of this paper is thus to modulate Wnt activation and quantify the response of entire *Hydra* tissue spheres in terms of regeneration, rheology and actin nematic order. To achieve this, we used two drugs, alsterpaullone and iCRT14, which were previously used in *Hydra* to demonstrate the involvement of the canonical Wnt pathway in axis definition (6, 22–26).

We achieved Wnt pathway activation by treating the animals with alsterpaullone (AP). It was previously demonstrated that treating full *Hydras* with AP induces the formation of ectopic tentacles along the body column (6). AP is a kinase inhibitor that specifically targets GSK3 (27). From a molecular biology perspective, GSK3 is a kinase that phosphorylates β-catenin. This phosphorylation inhibits β-catenin’s proliferation. When the Wnt pathway is activated, phosphorylation is blocked, allowing β-catenin to accumulate, translocate to the nucleus, and express target genes. AP therefore blocks this phosphorylation and forces the activation of the Wnt pathway even in the absence of ligands. A common protocol, described in (28), involves immersing full *Hydras* in 5µ𝑀 AP for 48ℎ, followed by transferring them back to normal conditions. Two to three days after the end of the treatment, the *Hydra* were puffy, with numerous small tentacles sprouting from their bodies. AP treatment gave head-like characteristics to the body column, including strong expression of both head organizer and head inhibitory signals (6). This explains the emergence of multiples tentacles spots without the formation of complete heads.

On the opposite, inhibition of the Wnt pathway was achieved by treating *Hydras* with iCRT14. It has been demonstrated (29) that this molecule specifically interferes between β-catenin and the transcription factor TCF4, thereby inhibiting the transcription of target genes in the Wnt pathway.

Previous works have shown that by inhibiting this signaling pathway, iCRT14 blocked *Hydra* regeneration after amputation (24, 25). Indeed, 72ℎ after bisection, *Hydras* treated with this drug, although they had closed the wound, exhibited a rounded oral end without any tentacle or hypostome formation (24). The same observation was made for the foot, where after 48ℎ, no cell differentiation had occurred (24).

In this paper, we treated full *Hydras* with AP or iCRT14 before excising tissue spheres which had a uniform high- or low-level activation of the Wnt pathway. Then, using large scale screening and timelapse imaging, our previously published microaspiration setup, transgenic *Hydra* lines and 3d spinning disk microscopy, we were able to quantify the effect of these two drugs on regeneration at the organism scale, tissue rheology, osmotic oscillations and actin nematic order. Overall, our results support the hypotheses of the two recent models of *Hydra* patterning. AP prevented full regeneration, induced clear defects in both patterning and morphogenesis which we explain by a softening of the tissues as well as a misalignment of myonemes. iCRT14, on the other hand, only partially blocked regeneration, showed no significant effect on tissue stiffness but induced a disassembly of actin filament. As a result, we propose in the Discussion section that the two current models might not be contradictory and that it would be interesting to merge them and test the predictions of the resulting framework. Having two different, albeit most likely coupled, mechanisms for axial patterning should add robustness to the system. Future work could then test whether such robustness could explain the great plasticity of *Hydra* regeneration from a variety of initial conditions.

## Methods

### *Hydra* care and lines used

*Hydras* were kept in a suitable environment called *Hydra* medium (HM) which consists of pure water (filtered with Direct-Q 3UV, Merck, Frankfurt, Germany), 0.5𝑚𝑀 𝑁_𝑎_𝐻𝐶𝑂_3_, 1𝑚𝑀 𝐶_𝑎_𝐶𝑙_2_, 0.1𝑚𝑀 𝑀𝑔𝐶𝑙_2_, 0.08𝑚𝑀 𝑀𝑔𝑆𝑂_4_and 0.03𝑚𝑀 𝐾𝑛𝑂_3_at a pH between 7 and 7.3 adjusted using 𝐻𝐶𝑙. The cultures were stored in Tupperware containers, which were then kept in an incubator (Pol-Ekko-Aparature, Wodzislaw Slaski, Poland) in the dark at a fixed temperature of 18℃. The *Hydras* were fed twice a week with newly hatched *Artemia* (Hobby; Dohse Aquaristik, Grafschaft, Germany). Three *Hydra vulgaris* lines were maintained and used for experiments: a wild type (AEP) line and transgenic strains. A reverse watermelon (RWM) line expressing GFP in the ectoderm and DsRed2 in the endoderm(RWM) (30), both under the control of an active actin promoter and an ectoderm LifeAct-GFP line (31). AEP samples were used for regeneration and circularity measurements, RWM for mechanical characterizations and measurements of the osmotic oscillations and ectoderm-LifeAct-GFP for the analysis of the actin structure.

### Preparation of tissue spheres and pharmacological treatments

Tissue spheres were made by hand-cutting slices of tissue from the body column of *Hydras* which had been starved for 24 to 48h, using a sterile scalpel (Bistouri UU no. 10, Holtex, Aix-en-Provence, France). First, the head and foot were removed via two transverse cuts. Then, we cut 3-4 slices from the remaining body column. This resulted in ring-shaped pieces. These were then cut in half to form strips which we left to close into a sphere for 3 to 4 hours. These cuts were made until 6-8 pieces were obtained from a single *Hydra*. To improve their precision, these manipulations were carried out under an illuminated stereomicroscope (Leica Microsystems, Wetzlar, Germany). After excision, all samples were left to fold back into tissue spheres for 3h. All experiments were started at this 3h post excision time point which we refer to as t_0_ throughout the manuscript.

For Wnt modulation, we prepared solutions of 5µ𝑀 and 2.5 µ𝑀 AP with 0.025% DMSO (Sigma-Aldrich, St Louis, USA) or 5µ𝑀 and 1µ𝑀 iCRT14 (Sigma-Aldrich) with 0.05% DMSO in HM. These solutions were prepared from stock solutions at 20𝑚𝑀 (100% DMSO) for AP and 10𝑚𝑀 (100% DMSO) for iCRT14. Parental animals were pre-incubated in the respective drugs in HM for 2h (iCRT14) or 48h (AP) prior to excision. Tissue fragments were excised and incubated in the same drug-containing HM during folding into tissue spheres and the subsequent experiments. Images of iCRT14-treated samples at days 6 were acquired under a stereomicroscope at 4x magnification. Experiments performed on full Hydras followed the same protocol but they were not cut into tissue spheres.

### Young’s modulus measurements

All micro-aspiration experiments were carried out as previously described in (16) and using RWM tissue spheres. Since the experiments lasted for a few tens of minutes after t_0_, they were performed at 5𝜇𝑀 concentrations for both drugs. For each batch of six tissue spheres, the samples were inserted in the microfluidic channel at t_0_. We then applied a series of steps of applied pressures. This was achieved by varying the height of an open syringe using hydrostatic pressure. Between each pressure step, the samples were released for 5min to avoid any hysteresis coming from the previous aspiration. Images of the aspirated samples were acquired on a Leica DMi6 inverted fluorescence microscope equipped with a 10x objective, imaging the ectoderm of the samples. For each pressure step, we measured the length 𝐿 of tissue inside the holes. We then computed the real aspirated length as:

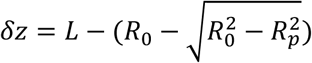

Where 𝑅_0_ is the radius of the tissue sphere and 𝑅_𝑝_ the radius of the hole, here 50μm.

Theoretically, we used the Saint-Venant Kirchhoff model and finite element simulations to show that in a certain applied pressure range (400 Pa – 4kPa), the length of aspirated tissue 𝛿𝑧 increases linearly with the applied pressure Δ𝑃 and that the Young’s modulus 𝐸 of the sample can be extracted using the following relationship (16):

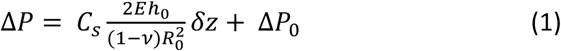

Where 𝐶_𝑠_ is a dimensionless pre-factor, 𝜈 is the Poisson coefficient, ℎ_0_and 𝑅_0_are its thickness and radius, respectively, and Δ𝑃_0_ is a negative intercept, signature of the non-linear elastic behavior at low applied pressures (16). We used this equation with ℎ_0_ = 20𝜇𝑚, 𝜈 = 0.495 and 𝐶_𝑆_ = 23.4 along with measurements of 𝑅_0_to extract the values of 𝐸.

The data shown in Fig 2d regarding the small deformation regime were obtained in the same experimental manner but with a lower pressure range of 50-1600 Pa. The curves represent the averages of n=22 for control samples and n=27 for AP-treated samples.

**Figure 1.**
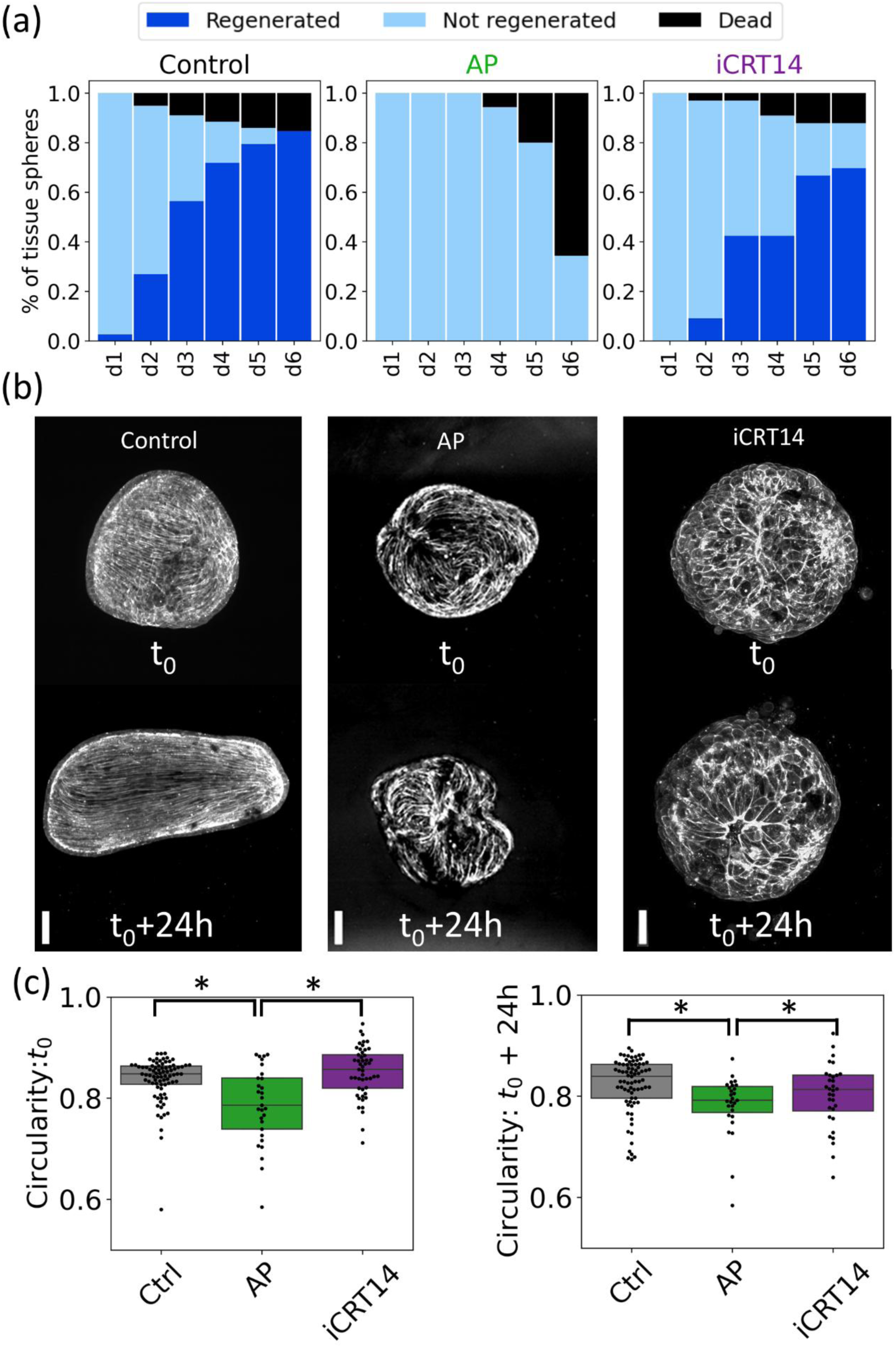
Effect of Wnt modulation on regeneration and morphogenesis of AEP Hydra tissue spheres. (a) Fraction of regenerated (dark blue), unregenerated (light blue) and dead (black) AEP Hydra tissue spheres from day 1 (d1) to day 6 (d6) after t_0_. Data is shown for controls on the left, AP-treated samples in the center and iCRT-treated samples on the right, n= 95,35 and 35, respectively. (b) Maximum intensity projections of LifeAct-GFP tissue spheres in control (left), AP-treated (center) and iCRT14-treated samples (right) at t_0_ and t_0_+24h, scale bars: 100 μm. (c) Quantification of circularity for all three treatments at t_0_ (n=82,29 and 49 in the order shown on the plot) and t_0_+24h (n=77,27 and 32). The boxes display the median and quartile of each distribution and stars represent statistical significance with a p-value<0.01.

**Figure 2.**
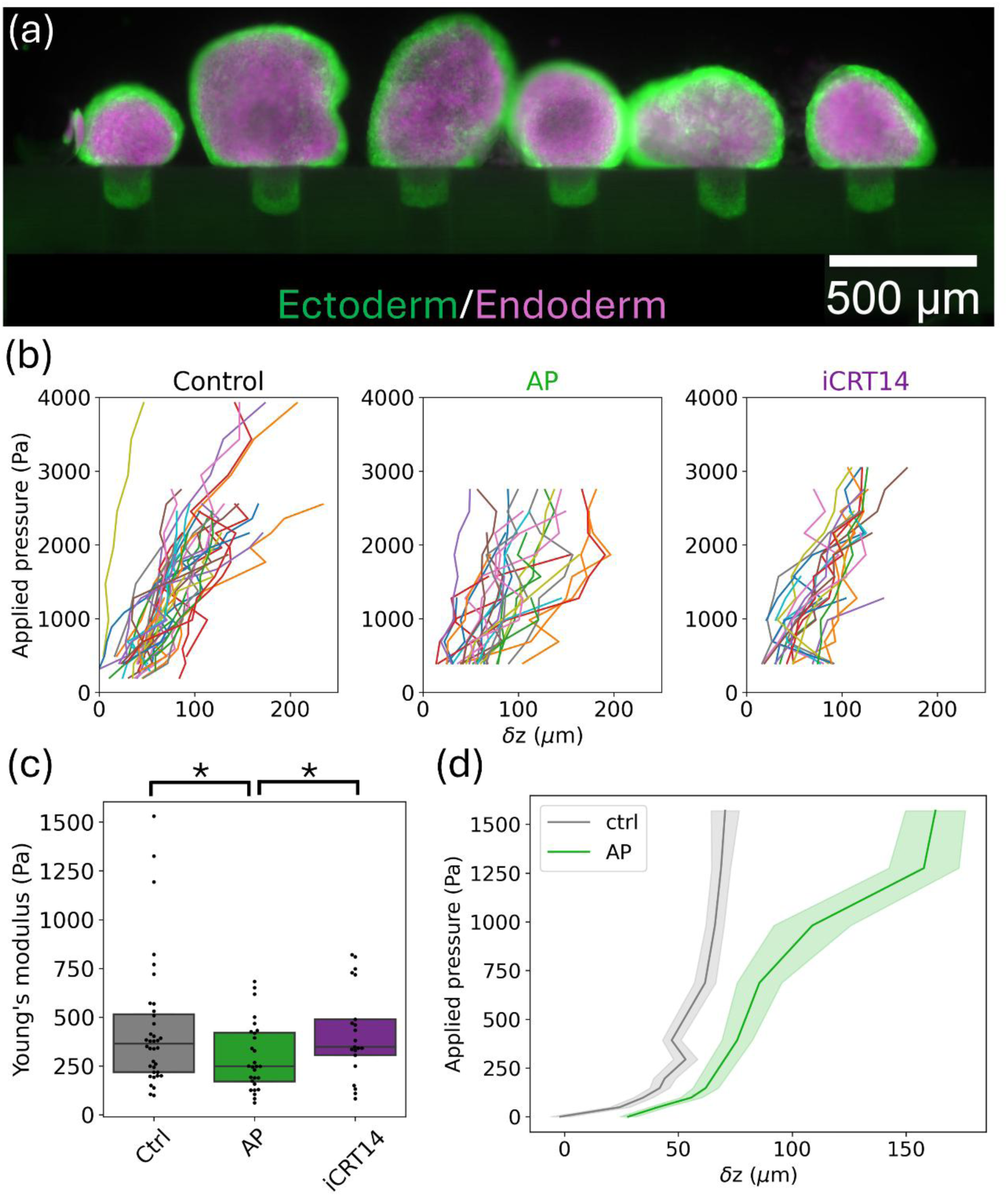
(a) Snapshot of six Hydra tissue spheres aspirated within the microfluidic device. (b) Plots of applied pressure versus aspirated length 𝛿𝑧 for controls on the left, AP-treated samples at 5μM in the center and iCRT-treated samples at 5μM on the right. Each colored line corresponds to a single sample, n= 36,28 and 21, respectively. (c) Distribution of Young’s moduli extracted from each sample shown (b) using linear fits and Equation 1. The boxes display the median and quartile of each distribution and stars represent statistical significance with a p-value<0.01.(d) Averaged responses of control (grey) and AP-treated samples (green) at low applied pressures. The lines represent the average and the shaded area the standard error of the mean, n=22 and 27.

Numerical simulations shown in Supplementary Figure 4 were also performed as described in (16) using Comsol Multiphysics (COMSOL Inc, Burlington, USA).

### Analysis of osmotic oscillations

For imaging, Reverse Watermelon (RWM) strain tissue spheres were individually placed in custom agarose wells. A 2% low melt agarose (Carl Roth, Karlsruhe, Germany) solution in HM was poured into a well of a 6-well plate (Thermo Fisher Scientific, Waltham, Massachusetts) to form a uniform layer approximately 5mm thick. After gel solidification, cylindrical holes of were punched into the agarose, using a 1mm Harris Uni-Core puncher (Sigma-Aldrich), to create individual imaging chambers. Each well was then filled with HM and covered with a thin layer of paraffin oil (VWR, Radnor, USA) to prevent evaporation. For experiments requiring both agarose wells and the application of AP or ICRT14 on *Hydra* tissue spheres, the drugs were added to the medium and to the HM used to make the gel to prevent dilution effects. Care was given to add the drugs in the agarose at the latest possible time to avoid heating effects on the stability of the drugs.

Time-lapse imaging was carried out over a continuous period of 24 h, with images acquired every 15 min on a Leica DMi6 inverted fluorescence microscope equipped with a 10x objective.

Image sequences from the RFP channel were batch processed using ImageJ. Initial segmentation was performed via Li’s thresholding method, followed by morphological operations: hole filling, erosion (to separate adjacent particles), and dilation (to recover eroded areas). Only regions with an area exceeding 10 000μm^2^ and circularity greater than 0.30 were retained for analysis. A custom-made macro automated this workflow. Assuming radial symmetry, the projected area 𝐴 was converted to radius 𝑅 using: 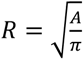. Phase I oscillations were identified manually based on amplitude and periodicity, this identification being performed blindly with respect to the experimental conditions. A Python script normalized radii to the minimum radius within each tracked series and detected peaks using the find_peaks function. For each oscillation, the following parameters were computed: duration (time between successive minima), maximal radius (peak value), and swelling rate (slope of a linear regression between the first minimum and the maximum radius). All oscillatory metrics were averaged per sphere for statistical comparisons.

### Quantification of regeneration

For each experiment, a total of 30 tissue spheres from AEP animals were prepared and kept in 90mm Petri dishes. They were visually assessed at t_0_ and every 24h after that for six days. Each sample was classified as either dead, regenerated as soon as a tentacle was identified and as alive but not regenerated otherwise. HM supplemented with the drugs was renewed every 48h.

For circularity measurements, we acquired images of these samples under an epifluorescence DMI6 microscope (Leica Microsystems) equipped with a 10x objective at t_0_ and t_0_+24h. The resulting images were binarized in ImageJ (NIH, Bethesda, USA) and the projected area 𝐴 and perimeter 𝑃 of the sample was extracted. Circularity, 𝐶, was then computed as

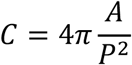

### Observation of actin nematic order

We used LifeAct-ectoderm GFP tissue spheres and full *Hydras*. Images were acquired as zstacks under an Eclipse Ti2 microscope (Nikon Europe, Amstelveen, Netherlans) equipped with a CSU-W1 spinning disk unit (Yokogawa, Tokyo, Japan) and an Orca Fusion BT camera (Hamamatsu, Shizuoka, Japan).

Optical planes were acquired every 2μm and maximum intensity projections were generated in ImageJ. The full 3d stacks were further treated to separate the basal and apical surfaces. To do so, we reproduced the routine published in (15) and available at https://github.com/KinneretLab. Briefly, this algorithm computes cost functions from the full 3d stacks by using an anisotropic diffusion filter in Matlab. Then, the two layers, basal and apical are extracted from the cost functions using the “Minimum Cost Z surface Projection” plugin in ImageJ. 2d projections of these curved surfaces are then generated from the height maps in Matlab.

We then used the basal projections and the areas with visible myonemes were manually drawn while the outline of each sample was automatically segmented in ImageJ from maximum intensity projections. We then computed the ratio of the surface covered by myonemes to the total surface of the sample.

### Statistical analysis

All plots were designed in Python. In all figures, the box plots represent the median and the quartiles of the corresponding distributions. Individual measurements are also shown as black dots. In most cases (circularity, Young’s modulus, duration of oscillations, maximum elongation, relative swelling rate, fraction of surface covered by myonemens), statistical tests were performed for each pair of conditions using a two-sample Student’s t-test. This was motivated by the apparent normal distribution of these measurements. For comparing the number of oscillations, however, we performed a Wilcoxon rank-sum test.

Regardless of the test used, we assumed statistical relevance if the p-value of this test was lower than 1% and this was represented as stars on the corresponding plots.

## Results

### Effect of Wnt modulation on *Hydra* regeneration

To study the effect of perturbing the canonical Wnt pathway on *Hydra* regeneration, we started by focusing on *Hydra* tissue spheres excised from organisms either untreated or pre-incubated with 5 μM AP for 48h or 5μM iCRT14 for 2h. These concentrations and time scales of exposure were previously used in the field but were not studied extensively on excised tissue spheres and their ensuing regeneration. Still, these specific treatments have been shown to have clear effects both at the molecular level (6, 25) and in terms of regeneration after bissection (22, 24, 32, 33).

After excision, samples were kept exposed to the drugs to ensure their effect would be maintained during further experiments. They were left to fold back into tissue spheres for three hours at which point the experiments were started at a time we refer to as t_0_.

The first effect of Wnt modulation we decided to study was on the outcome of regeneration. We followed tissue spheres for six days and we counted, each day, how many were dead, how many still resembled tissue spheres and how many were regenerated. Of note, to avoid biases, we counted a sample as regenerated as soon as it showed at least one tentacle, in line with previous work (13).

Unexpectedly, we found that the concentrations we used led to extensive lethality. In the first 24h, almost half of the iCRT14-treated samples died (Supplementary Figure 1). For AP, lethality occurred later but by day 3, all samples had disintegrated (Supplementary Figure 1). This is different from Wnt-overexpressing *Hydras* (5) which are viable, suggesting that AP had a lethal effect beyond its role as a Wnt activator. Overall, we found that these concentrations were not well suited for our study since we could not exclude potential non-specific effects of the drugs.

For this reason, we reduced these concentrations while keeping the time scales of application. We found that lowering them to 2.5 μM for AP and 1 μM for iCRT14 mitigated the lethality issue. As expected, we found that a vast majority (around 90%) of untreated samples properly regenerated and did so within a typical time of 2-4 days (Figure 1(a)). For AP-treated samples, we observed no tentacle appearance over the course of six days and started seeing disintegrating tissue spheres at days 5 and 6. In addition, we noticed a clear effect on morphogenesis during the first 24h of the experiments. Control samples, as expected, started spherical and gradually acquired an elongated, ellipsoid shape (Figure 1(b)). In contrast, AP-treated samples deviated from a spherical shape at t_0_, showing bulges that became more pronounced over time (Figure 1(b)). After 24h, they often showed multiple bulges and seemed swollen (shown from visualization of actin in an ectoderm LifeAct-GFP line in Figure 1(b)). To quantify this effect, we measured the circularity of the samples at t_0_ and t_0_+24h in all three conditions. The circularity *C*, sometimes called roundness, of a two-dimensional figure measures its deviation from a circle (for which C=1). It remains high for ellipses and falls as the figure becomes more tortuous. We indeed found that at t_0_, the circularity of AP-treated samples was significantly lower than that of control (Figure 1(c)). At longer time scales, we did not see major changes in their shapes and, in particular, never observed the appearance of tentacles and hypostomes. 24h later, the mean circularity had decreased for all conditions but remained lower for AP-treated samples (Figure 1(c)) than for the others. This suggests an important role of Wnt signaling in *Hydra* not only for patterning but also for morphogenesis.

iCRT14 treatments showed a mixed effect. Lethality was similar to controls, but only 23/35 tissue spheres still managed to regrow at least one tentacle. For these, we also observed a delay in tentacle appearance when compared to controls. Furthermore, of these 23, 7 showed a phenotype of improper regeneration where a protrusion resembling a tentacle was observed but the overall structure of the samples clearly differed from that of a full *Hydra* (Supplementary Figure 2). At t_0_+24h, we also noticed that they remained largely spherical unlike controls (Figure 1(b)), even though we didn’t find significant differences in circularity, likely because circles and ellipses both retain high circularity.

These results show that the experimental conditions we used are relevant. The observed effects on regeneration times and morphology confirm the known effects of AP and iCRT14 on Wnt signalling.

Interestingly, our treatment with iCRT14 did not completely abolish regeneration. This result might seem counterintuitive. It is however in line with (24) which showed that, even at 1 μM, regeneration after bisection was not fully blocked and that some samples still managed to regenerate either the head or foot. Only higher concentrations fully blocked regeneration. This points towards a partial inhibition of the canonical Wnt pathway and possible rescue mechanisms in patterning. Finally, the swollen aspects of AP-treated tissue spheres suggested changes in their mechanical properties which we decided to investigate next.

### Wnt activation softens *Hydra* tissue spheres

As described in the introduction, a recent model of *Hydra* patterning relies on the idea that Wnt activation could induce an increase in tissue deformability (19). The authors tested this hypothesis using a Wnt-overexpressing *Hydra* line and measured surface tensions using micro-aspiration.

Recently, we performed a thorough mechanical characterization of regenerating tissue spheres insisting on the importance of the non-linear elastic properties of these samples (16) and their behavior as 3d elastic shells rather than 2d surfaces. We therefore decided to repeat this characterization under both AP and iCRT14 treatments using RWM *Hydras* for visualization purposes.

Experimentally, we used a custom-made microfluidic device (Supplementary Figure 3) allowing to perform six micro-aspirations in parallel (Figure 2(a)) (16, 34). We have previously shown that, in a certain range of applied pressure Δ𝑃 (400 Pa – 4kPa), the length of aspirated tissue 𝛿𝑧 increased linearly with the applied pressure and that the Young’s modulus 𝐸 of the sample could be extracted using linear fits of this relationship (see Methods).

When we applied the same approach to AP and iCRT treated samples, we first observed that the linear relationship between applied pressure and aspirated length still held (Figure 2(b)). Using Equation 1, we then extracted measurements of 𝐸 for each sample based on the slope of the linear relationship and a precise measurement of 𝑅_0_ . Doing so, we found that AP significantly decreased the elastic modulus of *Hydra* tissue spheres (Figure 2(c)) from (4.4 ± 3.3).10^2^ Pa for control samples to (2.9 ± 1.7).10^2^ Pa. On the other hand, we found no difference between controls and iCRT14-treated samples (Figure 2(d)) which had an elastic modulus of (4.2 ± 2.2).10^2^ Pa. Images of aspirated tissue spheres in all three conditions but at the same applied pressure are shown in Supplementary figure 4.

Of note, we observed important sample-to-sample variability both in the values of 𝛿𝑧 and in the extracted Young’s moduli. We had previously observed this effect on non-treated sample and had found a weak, positive correlation between the sizes of the tissue spheres and their elasticity (16). Since these measurements were performed at t_0_, another potential source of variability is non-homogenous rheological properties. At this time, we do not expect patterning by Wnt to have any local effect yet. Instead, as the tissue spheres were newly reformed, it is possible that at the location of wound closure, the samples had a weak point, translating to a different elasticity. Our current setup, unfortunately, does not allow for a precise control of the orientation of the tissue spheres or of the location at which aspiration is performed making the study of local inhomogeneities a future interesting improvement from this work.

To measure 𝐸, we focused solely on the seemingly linear regime between applied pressures and deformations. This regime was predicted from numerical simulations of the Saint-Venant Kirchhoff model applied to spherical shells (16). This model also predicted the behavior of the tissue spheres in the regime of low applied stresses and small deformations. Until now, we had not validated that our samples indeed respected these predictions. To do so, we turned to a range of applied pressures between 50 and 1500 Pa and observed a non-linear relationship between the aspirated length and the applied pressure (Figure 2(d)). This offered a direct observation of a stress stiffening effect in *Hydra* tissue spheres. Of note, the AP-induced difference in stiffness was also clearly visible in this regime. For a given applied pressure, AP-treated samples were significantly more deformed than control ones (Figure 2(d)).

We then checked whether our measurements closely matched the predictions of the model in that small deformation regime. Since there is no analytical expression relating 𝛿𝑧 and Δ𝑃 in the geometry of micropipette aspiration in this regime, we reverted to numerical simulations. Using a finite element framework, we simulated the aspiration of an elastic spherical shell under the Saint-Venant Kirchoff rheology and with parameters matching those of the experiments. By trial and error, we modified the elastic modulus of the shell until we managed to reproduce measurements made on a single sample.

We found very good agreement for a value of 𝐸 = 750𝑃𝑎 (Supplementary Figure 2), validating the use of this specific rheological model to describe *Hydra* tissue spheres.

Overall, these results strongly support the hypothesis of (19), namely that Wnt activation increases deformability, i.e. lowering stiffness. Although our experiments were performed on tissue spheres where Wnt was either activated or inhibited everywhere, it is reasonable to imagine that a local activation of the Wnt pathway, as it occurs in normal regeneration, would lead to a local reduction in elastic modulus at later time points. Then, under an isotropic influx of water due to osmotic differences, it is expected that a spherical shell of anisotropic stiffness will deform more where it is softer and less where it is stiffer, providing a clear mechanical basis for the strain localization at the heart of the model of (19). On the other hand, the lack of impact of iCRT14 was to be expected.

Inhibition of Wnt activation should only induce small, local differences with control samples on which Wnt activation is localized. Therefore, iCRT14-treated samples should be mechanically very similar to untreated ones. Finally, our data clearly demonstrate the non-linear nature of the elastic response of *Hydra* tissue spheres (Figure 2(d)) which was only indirectly observed in (16). We argue that this nonlinear elastic response could be important in *Hydra* patterning.

A natural consequence of these changes in mechanical properties is the effect they might have on the osmotic oscillations observed during *Hydra* regeneration which are known to play a role in patterning (11, 13).

### Effect of Wnt modulation on osmotic oscillations

To characterize osmotic oscillations, we used a transgenic *Hydra* line expressing RFP in the ectoderm (30). We placed, at t_0_, up to 40 tissue spheres in individual microwells made in agarose gels, acquired high resolution time lapses for each of them and extracted their radius over time 𝑅(𝑡).

We then decided to focus on the so-called Phase I oscillations which are the relevant ones for *Hydra* patterning (11, 12). We manually selected these specific oscillations as the earliest, longest and with the largest amplitude and identifying the switch to Phase II oscillations (Figure 3(a)). In line with the results of (19), we observed a potential difference for AP-treated samples which seemed to display oscillations of higher amplitude and longer duration (Figure 3(a)). To better characterize these oscillations, we focused on four parameters (Supplementary Figure 6).

**Figure 3.**
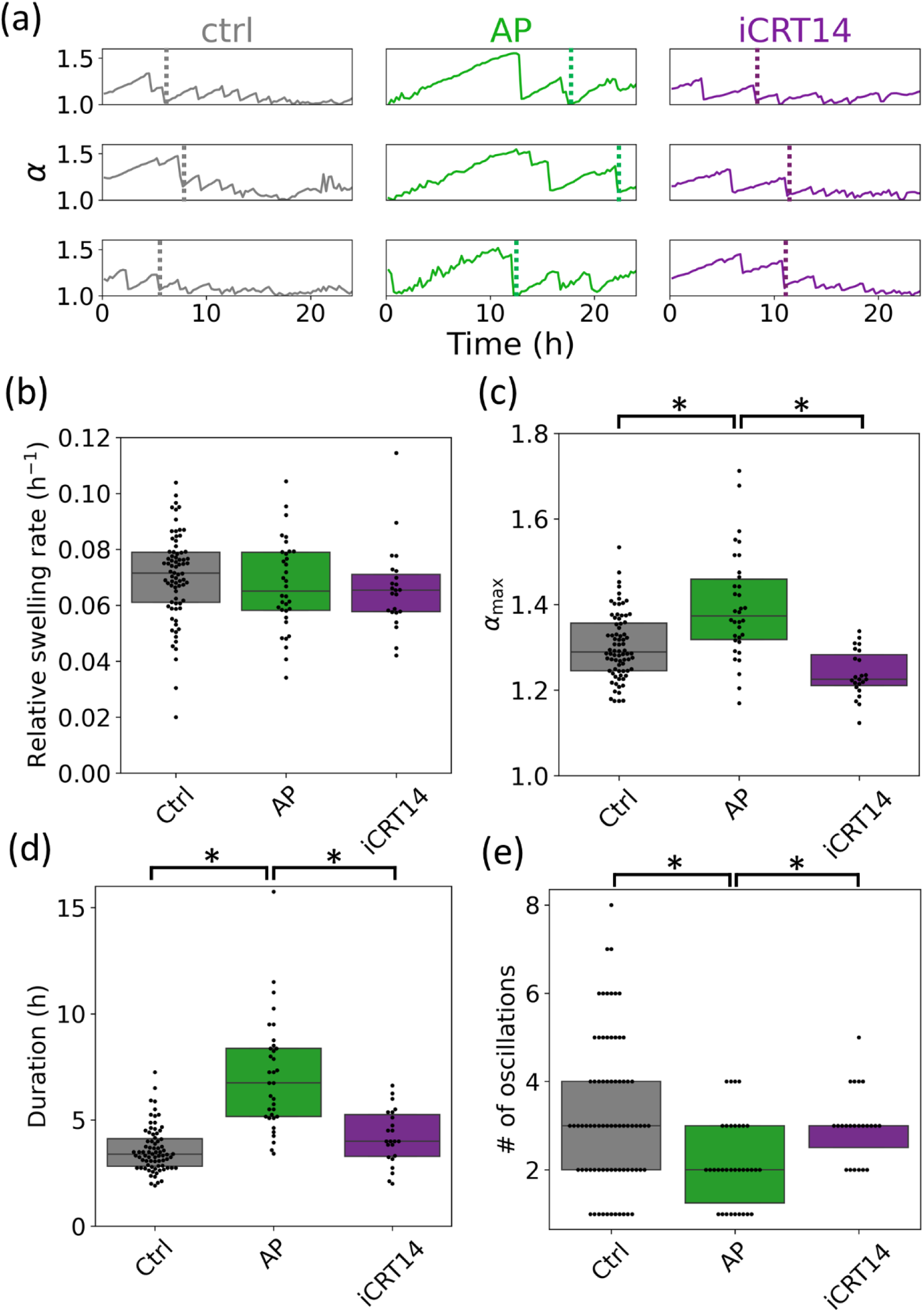
(a) Three examples of the dynamics of 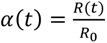 for each of the three different conditions, controls in grey, AP in green and iCRT in purple using RWM Hydras. The dashed color lines indicate the time of the manually detected switch from phase I to phase II oscillations. (b-e) quantification of all four parameters characterizing osmotic oscillations: relative swelling rate in (b), maximum stretch ratio in (c), duration of oscillations in (d) and number of phase I oscillations in (e). In each, boxes display the median and quartile of each distribution and stars represent statistical significance with a p-value<0.01. n=76,36 and 23, respectively.

First, the swelling rate 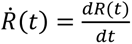 is the average speed at which the samples swell. Previous work has shown that this swelling rate increases proportionally with the rest radius 𝑅_0_ of the tissues spheres (11) and we report the relative swelling rate: 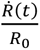. We did not find any differences in the relative swelling rate between control, AP-treated and iCRT14-treated samples (Figure 3(b)). Since this swelling is known to be due to osmotic differences between the inside and outside medium of the tissue spheres, we thus conclude that modulation of the Wnt pathway does not change either the osmolarity of the inner medium or the water permeability of the tissue spheres.

Second, the stretch ratio 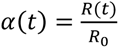 is a measure related to the mechanical strain 𝜀 of the spheres through 𝜀 = 𝛼 − 1. An important measurement is the maximum stretch ratio, 𝛼_𝑚𝑎𝑥_, at which the tissue spheres rupture. Indeed, since tissue rupture is due to a failure of cell-cell adhesions (16), it is thought to occur at a certain critical stress. The stresses experienced by the tissue increasing with both the strain and the stiffness of the tissue, a decrease in stiffness is expected to lead to an increase in the maximum stretch ratio. This is what we observed, with AP-treated samples showing a significant increase in 𝛼_𝑚𝑎𝑥_, from 1.30 ± 0.08 for controls to 1.41 ± 0.12 for AP-treated samples (Figure 3(c)). Of note, this corresponded to an increase in the maximum mechanical strain 𝜀_𝑚𝑎𝑥_ of around 30-40% which is of the same order as the observed changes in Young’s modulus between these two conditions. This would imply that the critical stress at which the tissues rupture are similar. Finally, as expected, we did not find any significant difference in 𝛼_𝑚𝑎𝑥_ for iCRT-treated samples for which 𝛼_𝑚𝑎𝑥_ was 1.25 ± 0.05 (Figure 3(c)).

Third, we measured the average duration of osmotic oscillations in each condition. Since we showed the relative swelling rate to be constant, we expected the average duration of oscillations to directly correlate with 𝛼_𝑚𝑎𝑥_. We found the average durations to be 3.61 ± 1.05h, 6.98 ± 2.55h and 4.20 ± 1.25h for controls, AP-treated and iCRT-treated samples, respectively (Figure 3(d)).

Finally, we counted the number of Phase I oscillations before the switch to Phase II. In (13), the authors proposed that the key parameter controlling pattern formation and the switch to Phase II oscillations is the overall amount of tissue stretching, akin to the area under the curve of the radius overt time. If so, AP-treated samples should display fewer oscillations since each of these would be longer and larger. We were able to confirm this as we observed a significant decrease in the number of oscillations for AP-treated samples but not for iCRT14-treated ones (Figure 3(e)).

All these results are an indirect confirmation of a softening of the tissue spheres when the canonical Wnt pathway is activated. The applied drugs, on the other hand, had no effect on the swelling rates, indicating that swelling is fully controlled by osmotic fluxes and not mechanical properties. These results are also in line with those of (19) which were obtained with a treatment of AP at 5μM, despite the large lethality we observed as soon as day 2 after excision with this concentration.

### Effect on actin nematics

The last key ingredient in *Hydra* patterning we investigated was the organization of ectodermal myonemes. To do so, we used a transgenic *Hydra* line expressing LifeAct-GFP in the ectoderm. This line allows an observation of both cortical actin, outlining the shape of the cells, and the myonemes. We used 3d spinning-disk microscopy to achieve high spatial resolution and separated the basal and apical surfaces of the ectodermal cells (15) to create 2d projections specifically of the basal side where the myonemes lay (see Methods and Supplementary Figure 7).

Our first observation was a modulation of the amount of well-aligned myonemes by both AP and iCRT14. We decided to quantify the amount of myonemes by measuring the fraction of the projected area of the samples they covered (Supplementary Figure 8). Of note, this area represents only half of the samples, the one facing the objective, as we were not able to image the tissue spheres from both sides in a controlled way.

In controls, confirming the results of (14), we observed that the actin nematic order was partially inherited from the parent *Hydra*. The fraction of area covered by aligned myonemes was of 44.7 ± 18.3% (mean ± standard deviation, Figure 4(a)). AP-treated samples showed a much larger coverage of myonemes with a surface fraction of 77.9 ± 6.9% while the opposite was true for iCRT14-treated samples which showed little to no well-aligned myonemes with a surface fraction covered of only 15.3 ± 13.7% (Figure 4(a)).

**Figure 4.**
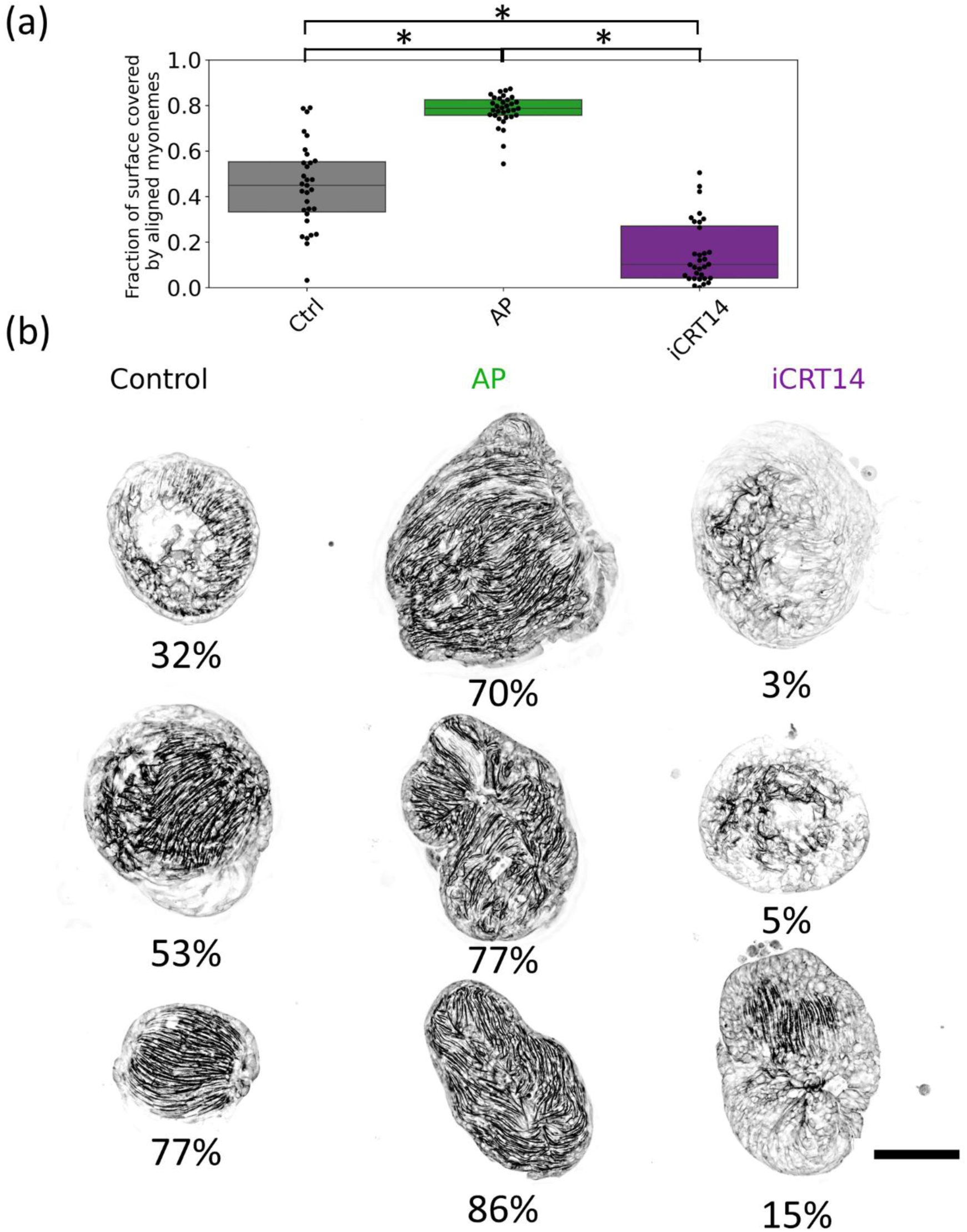
(a) Distributions of the fraction of their surfaces covered by visible myonemes in each condition with, controls on the left, AP-treated samples in the center and iCRT-treated samples on the right. n=31,33 and 32, respectively. In each, boxes display the median and quartile of each distribution and stars represent statistical significance with a p-value<0.01. (b) Three representative examples of basal surface projections of LifeAct-GFP Hydra tissue pieces in each condition shown in each column. The value shown below each image corresponds to its associated measurement of the surface fraction covered by visible myonemes. Scale bar: 200 microns.

In addition, we observed a change in the nematic order of these myonemes. In controls, they were clearly aligned along a single direction (Figure 4(b)) defining the future oral-aboral axis. In the iCRT14-treated samples which showed some significant myonemes, they also seemed to organize along a single axis (bottom example in Figure 4(b)). This partial, but normally organized, actin structure could explain the results of long-term regeneration experiments shown in Figure 1(a). Even under partial inhibition of the Wnt pathway, the nematic order could, when present, still induce local Wnt activation and proper regeneration.

The situation was strikingly different for AP-treated samples. Not only were the tissue spheres deformed but the actin nematic order seemed clearly perturbed and actin fibers appeared much wigglier and randomly aligned (Figure 4(b)). More examples of all three conditions supporting these observations can be found in Supplementary Figure 7.

Since myoneme organization is inherited, we then asked what the effect of Wnt modulation on the ectodermal myonemes of full *Hydras* was. As expected, controls showed a full coverage of myonemes and a perfect alignment in the oral-aboral axis (Supplementary Figure 9). For AP-treated full *Hydra*, we observed an altered organization. The filaments were wavy and curved (Supplementary Figure 9).

The shape of the *Hydras* seemed to follow this trend, being abnormally arched. The organization of fibers in iCRT14-treated whole *Hydras* was preserved and still showed a clear alignment along the head-tail axis 24ℎ after immersion in the drug. However, we observed in many *Hydras* the appearance of a ’cluster’ effect in the organization of cortical actin. The cell actin cortex became particularly visible in certain regions of the animal while myonemes were no longer visible in these regions.

Overall, these observations were similar to those in tissue spheres, and both can be summarized as follows: iCRT14 induces the disassembly of myonemes which still maintain their alignment when they are visible; AP stabilizes and enhances the formation of myonemes but disrupts their alignment which becomes multidirectional and seems to reduce the amount of cortical actin. This is in support of the hypothesis of (17) that actin nematic order aligns with gradients of Wnt activation. In the absence of Wnt activation, these myonemes slowly disappear whereas under strong, homogenous activation, they are numerous but do not align.

## Discussion

In this work, we used pharmacological means to modulate the activation of the Wnt pathway in a model organism for regeneration and axial patterning. By making the molecular landscape of *Hydra* tissue spheres homogenous, this approach allowed us to directly test different couplings between Wnt signalling and key ingredients in patterning: osmotic oscillations, rheology, actin nematic order.

In addition, we tested the effect of these drugs on the outcome of regeneration which, to our knowledge, had not been systematically assessed in this context.

Doing so, we observed and better characterized the non-linear elastic behavior of *Hydra* tissue spheres (Figure 2(d)). Of note, we showed that their elastic behavior held up to deformations as large as 30-40% before rupturing. This is uncommon but was observed for suspended MDCK monolayers (35) which also displayed a non-linear elastic rheology. In this system, the maximum deformations were even as large as 60%. Although *Hydra* tissue spheres are composed of two epithelial layers stuck together by extra-cellular matrix, their similarity with flat, suspended monolayers suggest that their elastic rheology might be a common signature of epithelia. How these biological systems can sustain such large deformations in an elastic regime remains unanswered and their response to deformations at the cellular scale should be investigated to better understand this ability.

Furthermore, we demonstrated that activation of the Wnt pathway has a direct impact on elasticity, making the tissues softer. Wnt, through the canonical and non-canonical pathways, has a wide array of effects at the cellular and tissular level linked to mechanical properties. For example, activation of the canonical pathway was shown to destabilize adherens junctions in cancer cell lines (36). One key mechanism is the recruitment of β-catenin from cadherin-based cell-cell junctions to the cytoplasm and nucleus, increasing the amount of free β-catenin and down-regulating genes involved in these junctions (37). Another effect of this pathway is to induce the endocytosis of focal adhesions (38).

Since these adhesions couple the cytoskeleton to the extra-cellular matrix, their endocytosis reduces the cell-matrix adhesion. These effects are not limited to the canonical Wnt pathway. For example, activation of the Wnt-PCP pathway in *Xenopus* embryos impacts the fibronectin matrix, Rac signaling and tissue tension (39). Which molecular and cellular mechanisms are involved in the observed softening of *Hydra* tissue spheres under Wnt activation is a question naturally occurring from our work.

We argue that this feedback between Wnt and elasticity has a particular significance in *Hydra*. Wnt activation has always been thought of as a patterning mechanism in this organism. The formation of a local hotspot of Wnt activation is seen as the first step of axis definition. While this remains true, our results suggest that this patterning by Wnt also induces heterogeneities in mechanical properties.

Such heterogeneities are known to be critical for morphogenesis: the development of the proper shape of organisms (40). Morphogenesis results from the interplay between the mechanical properties of the tissues and the forces acting on them. In other systems, it has been shown that both force and stiffness heterogeneities can lead to axis elongation (41, 42). They can direct both the direction of elongation and its dynamics. In the case of *Hydra*, one such driving force is osmotic swelling which is homogenous. Since mechanical properties are now thought to correlate with Wnt activity, one expects elongation of the samples in the direction of the Wnt hotspot. Therefore, the formation of this Wnt hotspot would be key both in the formation of the head organizer and in the morphogenetic event of axis elongation. This is further supported by our observation that AP induces defects in morphogenesis with the appearance of multiple bulges on regenerating tissue spheres (Figure 1(b-c)). Having molecular patterning and axis elongation controlled by a single system could provide plasticity which is a signature property of *Hydra* regeneration which can succeed from a variety of initial conditions.

A consequence of these changes in mechanical properties is the changes we observed in osmotic oscillations. Many biological structures share the lumen-like structure of *Hydra* tissue spheres: cysts in general which are a common motif in the brain, the heart, the gut etc … It has been shown in blastocysts that hydraulic oscillations, similar to osmotic oscillations in *Hydra*, play a role in controlling embryo size and stem cell fate (43). Theoretical work even proposed a framework explaining how these oscillations can regulate the size of growing luminal structures (44). This process could thus be quite general and applicable to a large variety of systems. Our results could provide a benchmark against which to test models further, offering perturbed conditions under which both the oscillations and the outcome of morphogenesis are impaired in a quantifiable way.

Another important finding of our work is the effect of Wnt modulation on actin organization. We found two major effects. First, Wnt activity seemed to correlate with the number of visible myonemes: the more Wnt activity, the more myonemes. Second, homogenous Wnt activation seemed to modify the alignment of these actin fibers. They were aligned in a single direction in control conditions and along multiple directions under AP treatment. These results are similar to some obtained in mammary epithelial cells. Using soluble Wnt3a proteins, the authors showed a stabilization and enhanced directional alignment of actin fibers (45). Similarly, (46) showed on hematopoietic stem cells that the Wnt ligand Wnt5a induces actin polymerization, increasing the amount of filamentous actin through a regulation of ROCK. Although the precise mechanisms at play in *Hydra* are still unknown, it has already been shown theoretically that actin alignment along a morphogen gradient could explain not only axial patterning but also budding, tentacle formation and the structure of the mouth (18). A coupling between nematic fields and diffusing chemical signals might thus offer a new, important class of patterning mechanisms which should be further studied both experimentally and theoretically. The ubiquity of the Wnt pathway in development and axis definition and its potential effects on both tissue rheology and actin organization strongly suggest its importance in the morphogenesis of many different organisms.

Finally, in the specific context of *Hydra* regeneration, our results should help settle ongoing debates. As described previously, the current state of the art proposes two different models, one focusing on actin nematic order (17) and the other on osmotic oscillations, changes in rheology and mechanically induced Wnt activation (19). Interestingly, our work supports the hypotheses underlying both models.

We therefore propose that they might not be mutually exclusive and that efforts should be made to try and combine them. Such a combination could be summarized as follows: either by random local fluctuations in mechanical properties, the existence of a partial actin nematic order or both, some regions of a regenerating tissue sphere will experience larger deformations. Because of the non-linear elasticity, such small fluctuations in deformations could be leveraged into larger fluctuations in stresses (16). These areas would then start expressing more Wnt ligands in accordance with (13). Based on our observations, Wnt activation would have a dual effet: a local softening of the tissue and a recruitment and alignment of myonemes. Softening would further localize deformations in these areas under the isotropic swelling due to the osmotic flux. If a +1 nematic defect pre-existed in that region, it would be stabilized, if not, it could form. As demonstrated in (15), such a +1 defect in a contractile network would further localize deformations and end up at the position of the head organizer. It would be particularly interesting to test whether such a mechanism would add redundancy to the patterning system and answer existing questions in the field. A key aspect which will need to be studied in the future is the dynamics of these different effects. In our work, we applied the drugs before running any experiments and did not explore how elasticity or actin organization evolved over time. Understanding the time required for Wnt signalling to feedback on these processes and comparing this time to that of regeneration should provide relevant information to fine tune the model we propose.

Importantly, we do not suggest that redundancy implies that the two processes are completely independent. In particular, it is well known that the rheology of epithelial tissues depends on both cortical tension in cells and on the strength of cell-cell adhesions (47). It is thus possible that the softening of the tissues observed under Wnt activation is a consequence of the changes in actin organization, potential changes in cortical actin and the number and alignment of myonemes. In turn, this actin organization could be re-organized, at least partially, by local levels of Wnt activation, as suggested by previous works on grafts of fully *Hydras* putting the two (Wnt gradient and myoneme orientation) at a right angle (33). It is also possible that local inhomogeneities in mechanical properties could feed back onto the actin structure.

Our point is that it would be interesting to try and encompass all the effects of Wnt modulation we described in a framework integrating them at the local scale. Another natural continuation of this work would be to entangle the molecular mechanisms responsible for the crosstalks between the different processes to understand causal relationships between them.

Our hope is that such an integrated model would help settle existing questions in the field. For example, one main weakness of the actin nematic framework was to explain how patterning occurred in cellular re-aggregates which do not inherit any myonemes. Notably, a recent study demonstrated that the emergence of a fully organized actin network still occurred in that situation, on the time scale of 100h (48). We argue that, in this situation, axial patterning could be mainly driven by osmotic oscillations and a feedback loop between tissue deformations, Wnt activation and tissue softening.

On the other hand, it was long thought that blocking osmotic oscillations by adding sucrose to the outer medium led to a complete inhibition of regeneration, all tissue spheres failing to break symmetry and define a new axis. Recent data has shown that although samples in isotonic conditions fail to regenerate, they still elongate along the axis of alignment of myonemes (17). In that situation, patterning and morphogenesis could be driven solely by acto-myosin contractility and tissue softening at the location of nematic defects driven by Wnt activation, even in the absence of osmotic oscillations. Another interesting question a new model could be tested on is the number of organizers emerging during *Hydra* regeneration. The Gierer-Meinhardt model, as a purely chemical reaction-diffusion one, has a characteristic wavelength, meaning that the number of hotspots of activator increases with system size. The mechano-chemical version, on the other hand, was shown numerically (19) and analytically (49) to stabilize a single organizer. Experimentally, excised tissue pieces consistently regenerate a single head. Cellular re-aggregates, however, can regrow multiple heads depending on their initial size. It would be interesting to test if the difference in the initial organization of the actin network, which is not taken into account in any of these models, could reconcile these different observations in an integrated framework. On the experimental side, it would be valuable to compare the predictions of such a model with the outcome of the regeneration of cellular aggregates in the absence of osmotic oscillations, a situation in which both drivers of patterning would be absent.

Further theoretical and numerical work will be needed to develop, simulate and confront the predictions of this model with all existing experimental observations. Still, we believe that the idea of a redundant system which could drive both patterning and morphogenesis remains exciting to understand the remarkable robustness and plasticity of *Hydra* regeneration.

## Conclusion

In this study, we pharmacologically modulated the canonical Wnt pathway in *Hydra* tissue spheres to investigate its impact on regeneration, tissue mechanics, and actin organization. We show that Wnt activation leads to tissue softening, disrupted actin alignment, altered osmotic oscillations, and impaired regeneration. Conversely, Wnt inhibition reduces actin filament density while preserving mechanical properties and partial regenerative capacity. These findings demonstrate that Wnt signaling affects not only axial patterning but also the mechanical and structural properties of regenerating tissues.

## Ackowledgments

This work was partially funded by the Mission pour les Initiatives Transverses et Interdisciplinaires of CNRS (Project MeChemReg to OCE and PM).

