## Supplementary Information for "Wnt signaling modulates tissue mechanics, actin order, and regeneration in *Hydra vulgaris*"

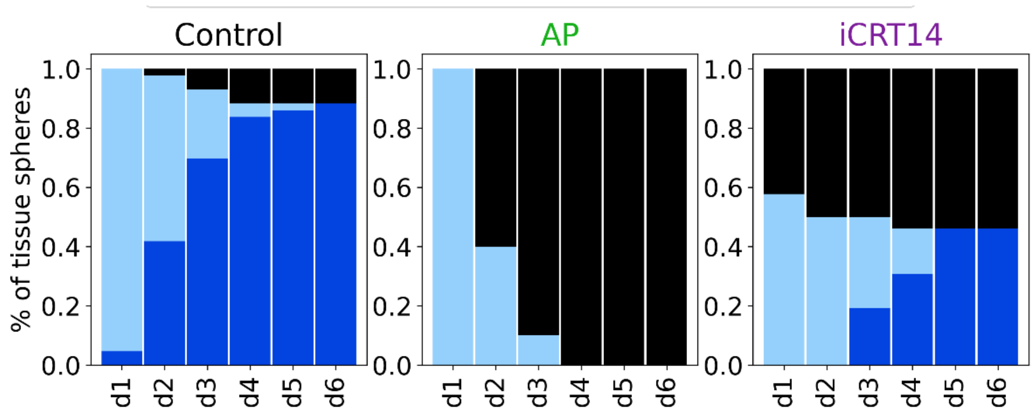


*Supplementary figure 1. Fraction of regenerated (dark blue), unregenerated (light blue) and dead (black) Hydra tissue spheres from day 1 (d1) to day 6 (d6) after $t_0$. Data is shown for controls on the left, AP-treated samples in the center and iCRT-treated samples on the right, n= 43,30 and 26, respectively.*

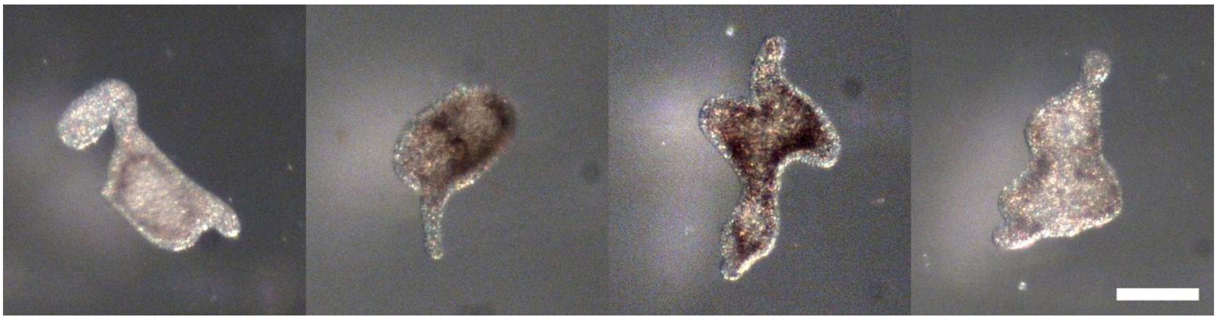

*Supplementary Figure 2. Examples of iCRT14-treated samples on day6 showing a tentacle but a clear defect in regeneration outcome. Scale bar: 2mm.*

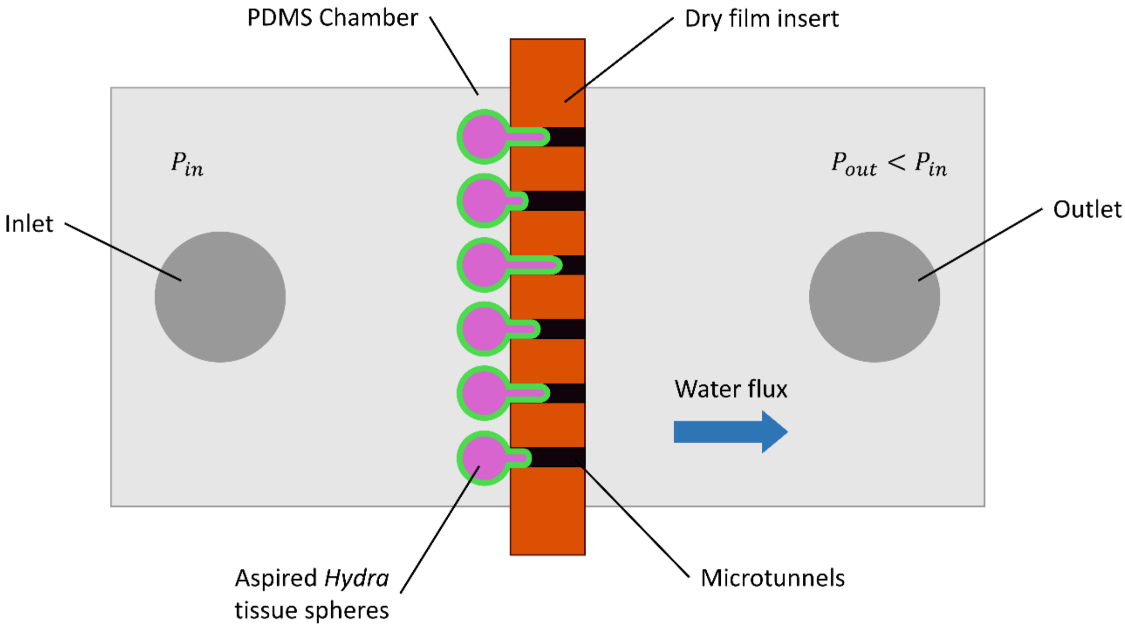


*Supplementary figure 3. Schematics of the parallelized micro-aspiration setup. A PDMS channel is barred by a dry film insert pierced with six cylindrical holes. A pressure difference is applied between the inlet and outlet through hydrostatic pressure. As a result, the tissue spheres (shown in green and magenta) are aspirated within the holes.*

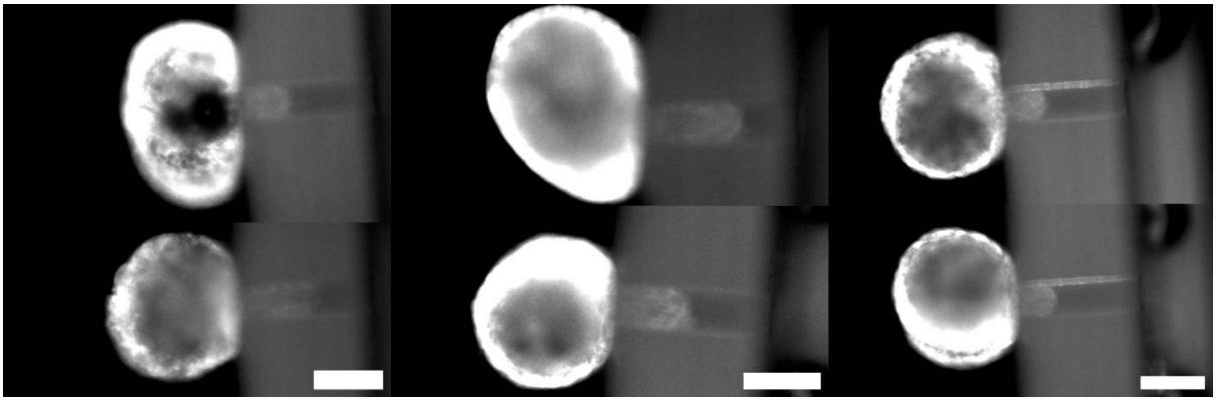

*Supplementary figure 4. Snapshots of tissue spheres aspirated in 100$\mu$m wide openings at $\Delta h = 16cm$. Controls on the left, AP-treated in the center and iCRT14-treated on the right. All scale bars are 200$\mu$m.*

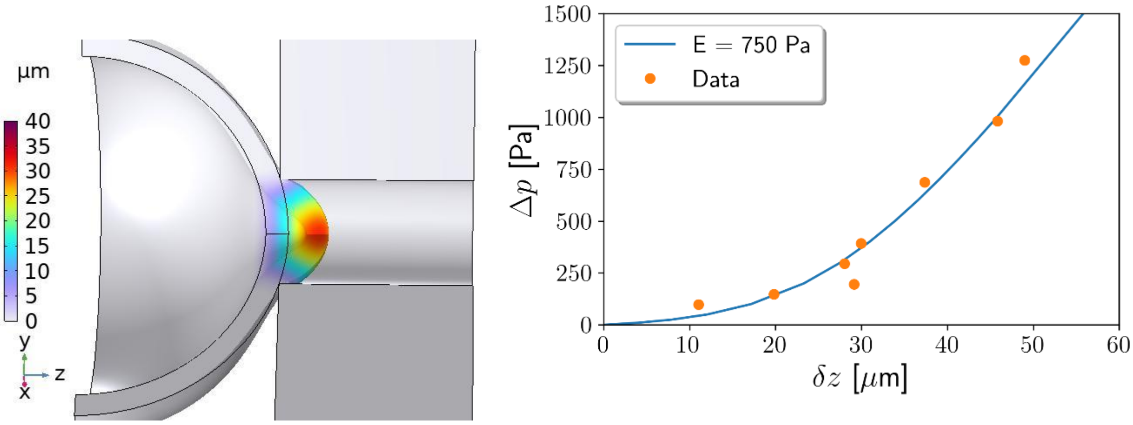


*Supplementary figure 5. Numerical simulations of low applied pressure regime. Left: snapshot of the finite element simulation. The color code represents the aspirated length. It is measured as a displacement fieldfrom the reference state of the spherical shell which is shown as black lines. Right: the blue line shows a stress-strain curve obtained from a simulation with E=750Pa. This value was adjusted by trial and error to match experimental measurements on a single, control sample, shown as orange dots.*

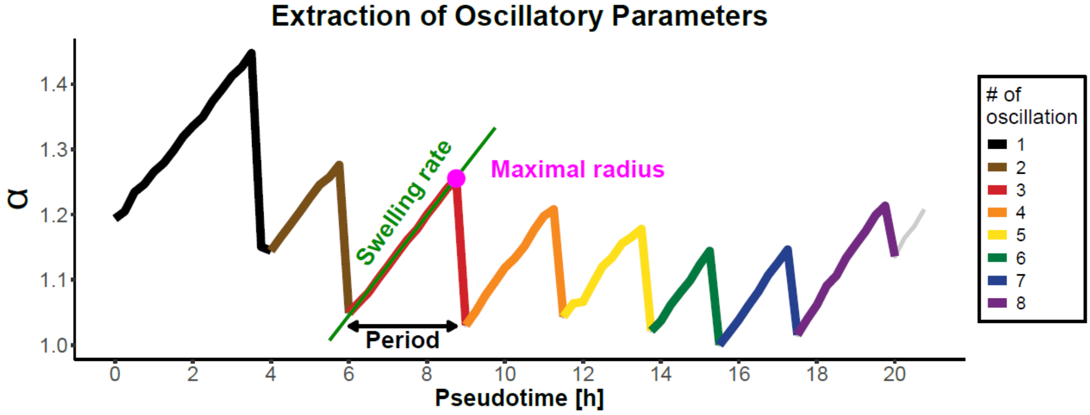


*Supplementary figure 6. Schematics of measured parameters for osmotic oscillations. Each oscillation, here with a total of 8, is shown in a different color.*

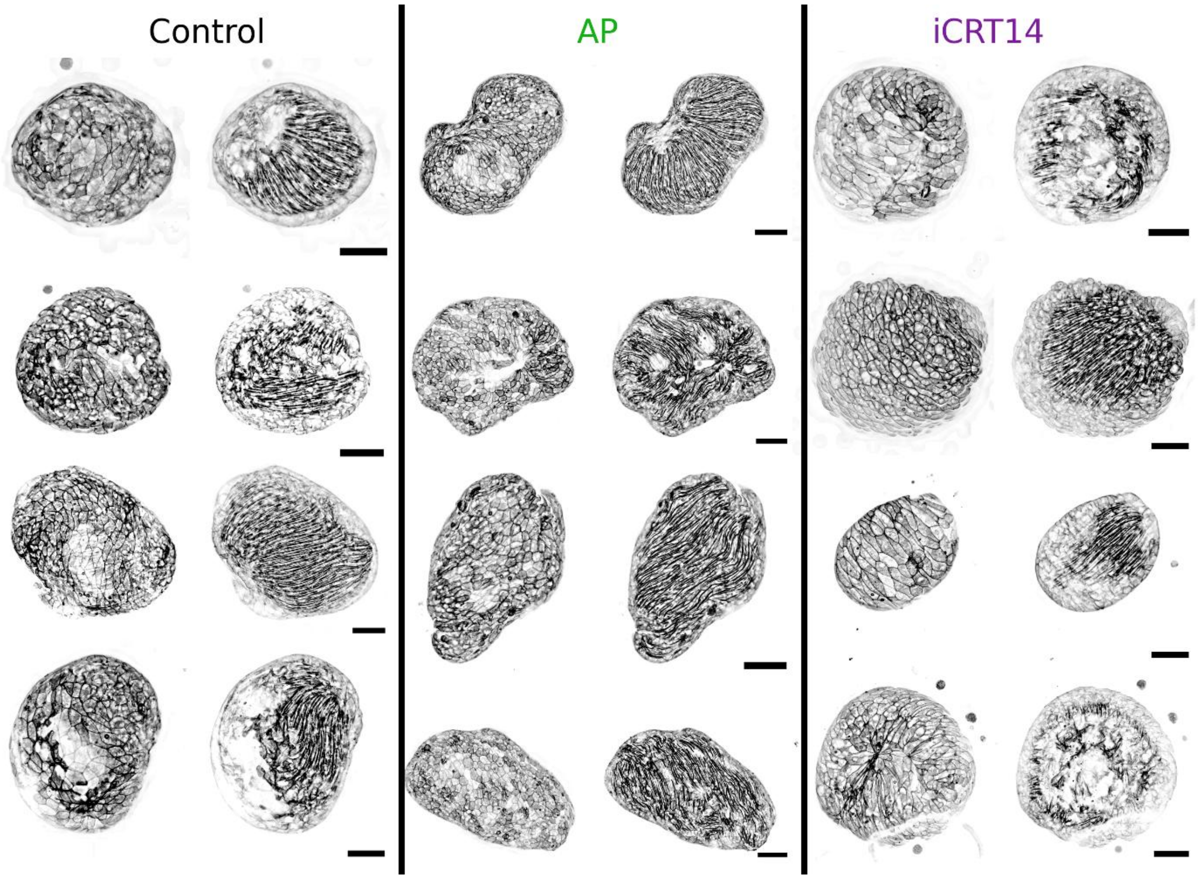


*Supplementary figure 7. Examples of surface separation in all three conditions. Four examples are given in each condition with, for each, the projection of the apical side showing cortical actin and cell shapes on the left, and the projection of the basal side, showing myonemes, on the right. All scale bars are 100 $\mu$m.*

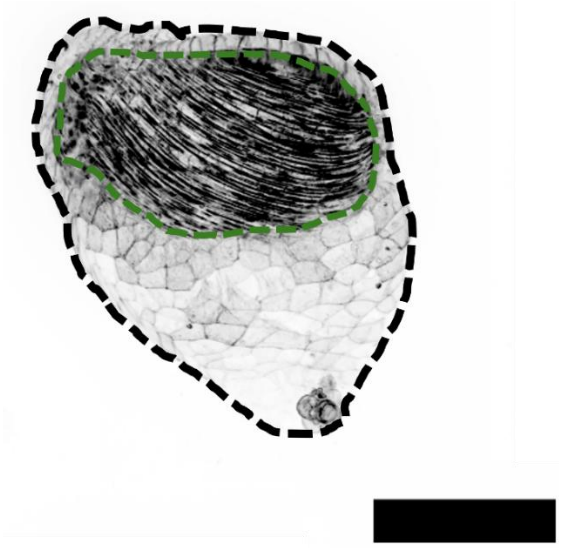

*Supplementary figure 8. Schematics of the quantification of the surface covered by myonemes. The raw image shows, in black, the maximum intensity projection of the fluorescence signal from a LifeAct-GFP tissue sphere. The dashed black line is the outline of the entire sample while the green dashed line is the outline of the area covered by visible myonemes. We measured the ratio of area of these two regions. Scale bar: 200 microns.*

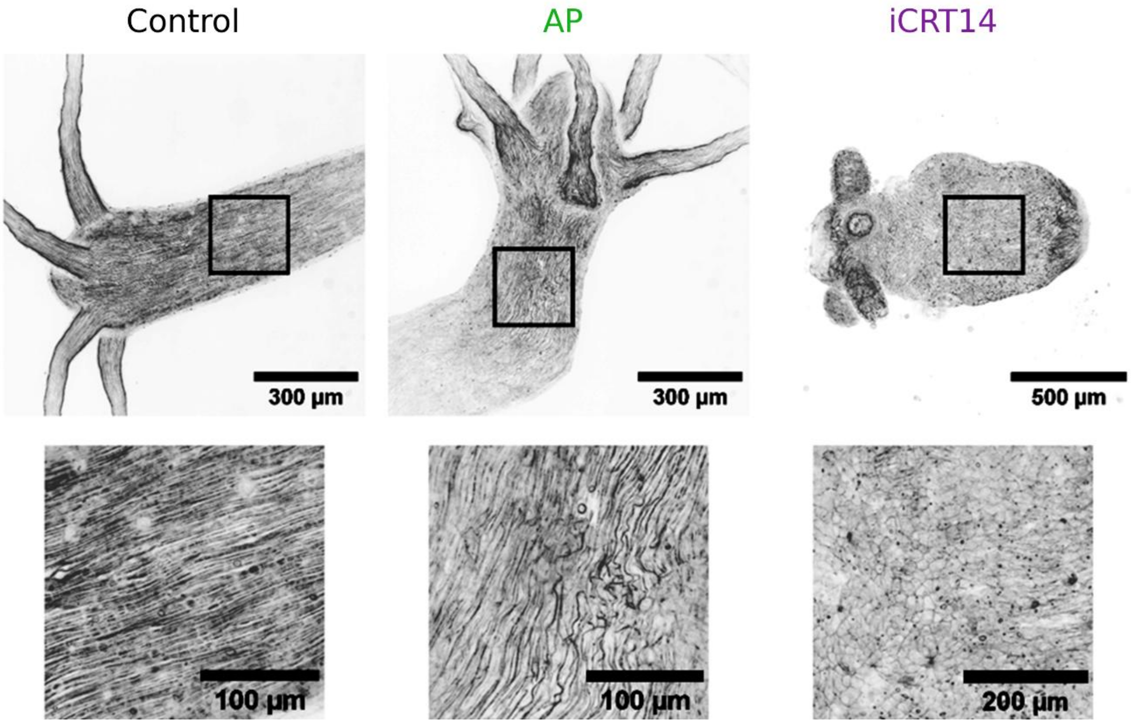


*Supplementary figure 9. Maximum intensity projections of LifeAct-GFP full Hydra in each condition at 5$\mu$M concentrations. The bottom row shows close ups of the regions delimited above by the black squares. Note the disorganization of myonemes under AP treatment and their local disappearance under iCRT14.*
